# Functional cooperativity mediated by rationally selected combinations of human monoclonal antibodies targeting the henipavirus receptor binding protein

**DOI:** 10.1101/2021.02.17.431743

**Authors:** Michael P. Doyle, Nurgun Kose, Viktoriya Borisevich, Elad Binshtein, Moushimi Amaya, Marcus Nagel, Edward J. Annand, Erica Armstrong, Robin Bombardi, Jinhui Dong, Kevin L. Schey, Christopher C. Broder, Larry Zeitlin, Erin A. Kuang, Zachary A. Bornholdt, Brandyn R. West, Thomas W. Geisbert, Robert W. Cross, James E. Crowe

**Author notes:** Corresponding author: James E. Crowe, Jr., MD, Mail: Vanderbilt Vaccine Center 11475 Medical Research Building IV 2213 Garland Avenue, Nashville, TN 37232-0417, USA.

## Abstract

Hendra virus (HeV) and Nipah virus (NiV), the prototypic members of the *Henipavirus* (HNV) genus, are emerging, zoonotic paramyxoviruses known to cause severe disease across six mammalian orders, including humans (Eaton et al., 2006). While several research groups have made strides in developing candidate vaccines and therapeutics against henipaviruses, such countermeasures have not been licensed for human use, and significant gaps in knowledge about the human immune response to these viruses exist. To address these gaps, we isolated a large panel of human monoclonal antibodies (mAbs) from the B cells of an individual with prior occupation-related exposure to the equine HeV vaccine (Equivac® HeV). Competition-binding and hydrogen-deuterium exchange mass spectrometry (HDX-MS) studies identified at least six distinct antigenic sites on the HeV/NiV receptor binding protein (RBP) that are recognized by human mAbs. Antibodies recognizing multiple antigenic sites potently neutralized NiV and/or HeV isolates *in vitro.* The most potent class of cross-reactive antibodies achieved neutralization by blocking viral attachment to the host cell receptors ephrin-B2 and ephrin-B3. Antibodies from this class mimic receptor binding by inducing a receptor-bound conformation to the HeV-RBP protein tetramer, exposing an epitope that appears to lie hidden in the interface between protomers within the HeV-RBP tetramer. Antibodies that recognize this cryptic epitope potently neutralized HeV and NiV. Flow cytometric studies using cell-surface-displayed HeV-RBP protein showed that cross-reactive, neutralizing mAbs from each of these classes cooperate for binding. In a highly stringent hamster model of NiV_B_ infection, antibodies from both classes reduced morbidity and mortality and achieved synergistic protection in combination and provided therapeutic benefit when combined into two bispecific platforms. These studies identified multiple candidate mAbs that might be suitable for use in a cocktail therapeutic approach to achieve synergistic antiviral potency and reduce the risk of virus escape during treatment.

## Introduction

Hendra virus (HeV) and Nipah virus (NiV), the prototypic henipaviruses, are emerging zoonotic paramyxoviruses known to cause severe disease in humans and diverse other mammalian orders. Multiple species of of *Pteropid* bats (flying foxes) act as reservoir hosts for these negative-sense, single-stranded RNA viruses in the *Paramyxoviridae* family with which they are understood to have co-evolved (Chua et al., 2002; Halpin et al., 2011; Halpin et al., 2000; Vidgen et al., 2015). HeV is transmitted from flying foxes to horses and from horses to in-contact humans causing severe respiratory and/or encephalitic disease mediated by endothelial vasculitis in both (Escaffre et al., 2013; Field, 2016; Murray et al., 1995a). HeV was identified in 1994, having caused the death of 14 of 21 infected horses and one of two infected humans in Queensland, Australia (Murray et al., 1995b; Selvey et al., 1995). Spillover has occurred sporadically with some seasonal and climatic trend since, causing disease in 105 horses and seven humans, with high case fatality rates (Queensland Government, 2020). NiV, which was discovered four years after HeV when hundreds of pig handlers fell ill with encephalitic disease (Chua et al., 1999), has continued to cause sporadic outbreaks in Bangladesh and India (Arunkumar et al., 2019; Soman Pillai et al., 2020). More direct routes of infection, including human-to-human transmission, and mortality rates approaching 100%, have been observed during recent NiV outbreaks (Chadha et al., 2006; Clayton et al., 2012; Gurley et al., 2007). Anthropogenic and climatic influences on flying foxes are affecting their roosting, feeding and migration habits as well as their susceptibility to heat-stress, disease and injury (Kessler et al., 2018; Plowright et al., 2015). These factors together with their resultant increase in intermediate host contact (humans and domestic animals) are associated with increasing geographic range and frequency of henipavirus disease spillover (Martin et al., 2018; Walsh et al., 2017). While HeV and NiV outbreaks historically have been confined geographically to Australia and Southeast Asia, respectively, risk of pandemic spread of these highly pathogenic agents related to regional and global population densities and difficulty avoiding international transmission via infected travelers has been highlighted by recent experience with SARS-CoV-2 (Morens and Fauci, 2020). Such consideration prompted the World Health Organization (WHO) to designate henipavirus infections as priority diseases requiring extensive and immediate research and development (Sweileh, 2017). The risk of global health crisis associated with henipaviruses is exacerbated by the lack of licensed antiviral drugs or vaccines for HNV and a dearth of knowledge of the human immune response to these viruses (Escaffre et al., 2013; Gomez Roman et al., 2020).

Passive immune transfer studies in both hamsters and ferrets have provided evidence that neutralizing antibodies are a correlate of immunoprotection from henipaviruses (Bossart et al., 2009; Guillaume et al., 2004; Guillaume et al., 2006). These data have been corroborated in multiple studies by investigators using murine, rabbit, or human antibody discovery technologies to isolate potently neutralizing antibodies to HeV and/or NiV (Aguilar et al., 2009; Mire et al., 2020; Zhu et al., 2006). One of these studies used phage display technology to isolate a human monoclonal antibody (hmAb), designated m102.4 (Zhu et al., 2008). This mAb potently neutralizes both HeV and NiV *in vitro* and protects against infection and disease in experimental henipavirus challenge models using ferrets or non-human primates (Bossart et al., 2011; Geisbert et al., 2014; Mire et al., 2016). More recently, two human mAbs, HENV-26 and HENV-32, were shown to neutralize HeV and NiV by distinct mechanisms and protect from NiV Bangladesh (NiV_B_) strain challenge in a ferret model (Dong et al., 2020). While these studies have laid a foundation for our understanding of how to target henipaviruses therapeutically, many questions remain regarding the antigenicity of the attachment glycoprotein, and whether escape mutations from these mAbs can develop *in vivo*.

Here, we isolated hmAbs from circulating B cells of an individual with occupation-related exposure to the equine HeV vaccine (Equivac^®^ HeV) (Middleton et al., 2014). Members of this large panel of antibodies target diverse antigenic sites, many of which are sites of vulnerability for neutralization for at least one virus. In particular, two functional classes of antibodies that we have termed “receptor-blocking” or “receptor-enhanced” neutralized HeV and NiV *in vitro* by distinct molecular mechanisms and provided protection when used as monotherapy against lethal challenge in hamsters with the highly virulent NiV_B_ strain. Antibodies recognizing these sites cooperate for binding to the henipavirus RBP glycoprotein that mediates attachment (formerly designated the G or glycosylated attachment protein, but recently renamed by the International Committee on Taxonomy of Viruses) (Rima et al., 2019). These mAbs also synergize for neutralization of both VSV-NiV_B_, as well as two Cedar viruses chimerized to display the HeV or NiV_B_ RBP and F surface glycoproteins. Cocktails of antibodies from these groups show superior therapeutic efficacy in hamsters, while bispecific antibodies bearing antigen binding fragments from both mAbs also show therapeutic benefit. In this model, “receptor-blocking” mAbs induce conformational changes to the RBP that better expose the “receptor-enhanced” antigenic site. These results suggest these mAbs could be used in a cocktail therapeutic approach to achieve synergistic neutralizing potency against henipavirus infections.

## RESULTS

### Cross-reactive, neutralizing antibodies target two distinct antigenic sites on HNV-RBP

Peripheral blood mononuclear cells (PBMCs) from an Australian veterinarian with occupation-related exposure to HeV-RBP (Equivac® HeV) were tested for secretion of antibodies binding to recombinant forms of the NiV attachment (RBP) glycoproteins for NiV_B_, the NiV Malaysian strain (NiV_M_), or HeV. In total, we isolated 41 distinct new mAbs that bind henipavirus RBPs. In order to group this large panel of mAbs rationally into those that recognized similar antigenic sites, we used a surface plasmon resonance platform to bin antibodies based on the antigenic sites they recognized on recombinant protein comprising the HeV-RBP head domain. This method immobilizes a first antibody on the surface of a gold-coated sensor-chip that captures soluble antigen, and then assesses the ability of a second antibody to bind to the captured antigen. The resulting data showed that mAbs binding to HeV-RBP recognized at least 6 distinct major antigenic sites, designated A-F (**Figure 1A, S1**).

**Figure 1.**
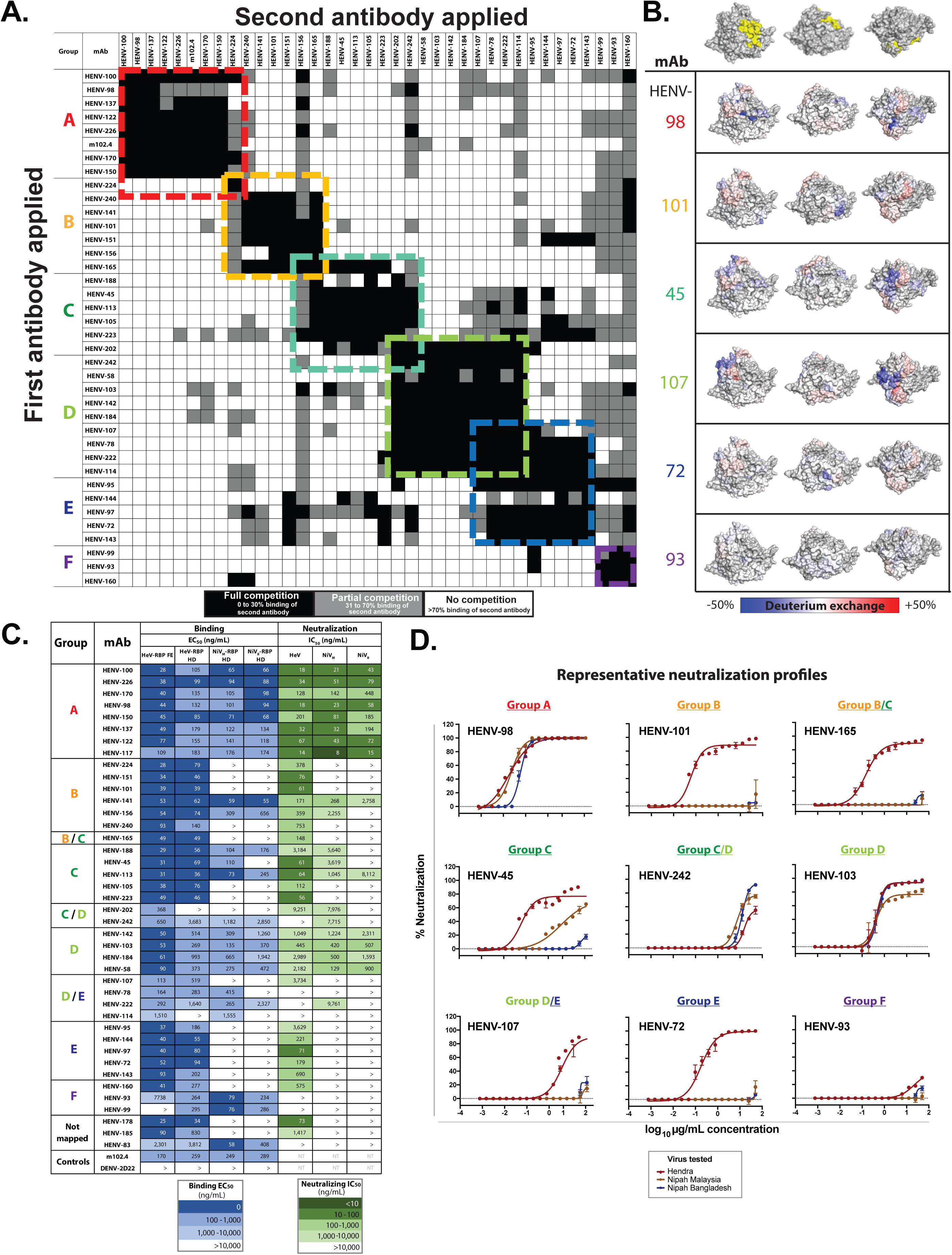
Identification of major antigenic sites for recognition of RBP by human mAbs. **A)** Surface plasmon resonance competition-binding of human antibodies against HeV-RBP. A first antibody was applied to a gold-coated sensorchip, and recombinant HeV-RBP head domain was associated to the coupled antibody. A second antibody was applied to the sensorchip to determine binding to RBP. Black boxes indicated a pairwise interaction in which the binding of the second antibody is blocked by the first. White indicates both antibodies could bind simultaneously. Gray indicates an intermediate competition phenotype. The matrix was assembled using the Carterra Epitope software. **B)** Hydrogen-deuterium exchange mass spectrometry profiles for representative mAbs. Decrease (blue) or increase (red) in deuterium exchange on HeV-RBP in the presence of antibody is mapped onto the crystal structure of HeV-RBP (PDB 6CMG). Structures are positioned in 3 orientations, with the top structure noting the ephrin-B2 binding site in yellow. **C)** Half maximal binding (blue) or neutralization (green) concentrations for antibodies against recombinant proteins or live HeV or NiV, respectively. **D)** Neutralization curve plots for representative antibodies against HeV, NiV Malaysia, or NiV Bangladesh viruses. Representative EC_50_ values for binding from 3 independent experiments are shown. IC_50_ values for neutralization are from a single independent experiment due to limitations of BSL-4 resources.

In tandem, we used hydrogen-deuterium exchange mass spectrometry (HDX-MS) to map the antigenic sites of representative antibodies from each group (**Figure 1B**), along with binding and neutralization assays to determine cross-reactivity and functional activity **(Figure 1C, D)**. Antibodies belonging to groups A and C cross-reacted with HeV, NiV_M_, and NiV_B_-RBP, and neutralized the corresponding viral strains. Group A, specifically, includes the control mAb m102.4, which has been thoroughly characterized for its ability to block viral attachment to the host cell receptors ephrin-B2 and ephrin-B3, and potently neutralize both HeV and NiV(2020; Xu et al., 2013) (Xu et al., 2013). As expected, a representative group A mAb HENV-98 caused a decrease in deuterium exchange in a region of the HeV-RBP that corresponds to the receptor-binding site. All group A mAbs also neutralized HeV, NiV_M_, and NiV_B_ strains *in vitro*. Notably, HENV-117 displayed exceptional potency, with half maximal inhibitory concentration (IC_50_) values of 14, 8, or 15 ng/mL against HeV, NiV_M_, or NiV_B_, respectively. To date, this is the most broad and potent neutralizing mAb targeting HeV and NiV ever described, suggesting it may possess superior therapeutic activity.

Group D represents a second class of mAbs that cross-neutralize HeV, NiV_M_, and NiV_B_, albeit with roughly 10-fold less potency than group A. The group D representative mAb HENV-107 mapped to a distinct site on the HeV-RPB head domain spanning the ß1 and ß6 propeller blades. This region of the head domain likely lies at the interface between protomers within the dimer-of-dimers structure of the HeV-RBP tetramer, suggesting a semi-cryptic site of vulnerability on RBP (Lee and Ataman, 2011). This region has been postulated to be important in fusion triggering, as point mutations made to this region render F unable to complete its fusion cascade (Aguilar et al., 2009; Liu et al., 2013).

While mAbs in group C display limited cross-neutralization of HeV and NiV, groups B and E contain mAbs that only neutralize HeV with appreciable potency. Group F mAbs are weakly neutralizing or non-neutralizing and appear to target an antigenic site that lies on the RBP face opposite the receptor-binding domain. This epitope is likely in a site that is poorly accessible in the membrane-anchored form of RBP, lending to the poor neutralizing activity observed for these mAbs. Overall, we discovered and mapped cross-reactive, neutralizing mAbs targeting two distinct major antigenic sites that likely use distinct mechanisms to achieve virus neutralization.

### Neutralizing mAbs either compete with, or are enhanced by, ephrin-B2 binding to HeV-RBP

With the knowledge that group A mAbs map to the receptor-binding domain of HeV-RBP, we sought to determine if these antibodies could block binding of soluble ephrin-B2 to cell surface-displayed HeV-RBP. 293F cells were transiently transfected with a cDNA construct encoding full-length HeV-RBP (head, stalk, transmembrane, and cytoplasmic domains) and incubated for 72 hours. These cells then were incubated with saturating concentrations of recombinant, soluble ephrin-B2, followed by addition of anti-RBP mAbs at a concentration of 2 µg/mL to assess the ability of antibodies to bind RPB in its receptor-bound state. Cells were analyzed by flow cytometry, comparing antibody binding in the presence or absence of ephrin-B2 (**Figure 2A)**. Antibodies in group A displayed a substantial decrease in binding in the presence of ephrin-B2, supporting the hypothesis that mAbs from this group potently neutralize by blocking binding of virus to host cells. This receptor-blocking phenotype is reflected in the activity of the control mAb m102.4, which also displayed decreased signal when associated to receptor-bound RBP.

**Figure 2.**
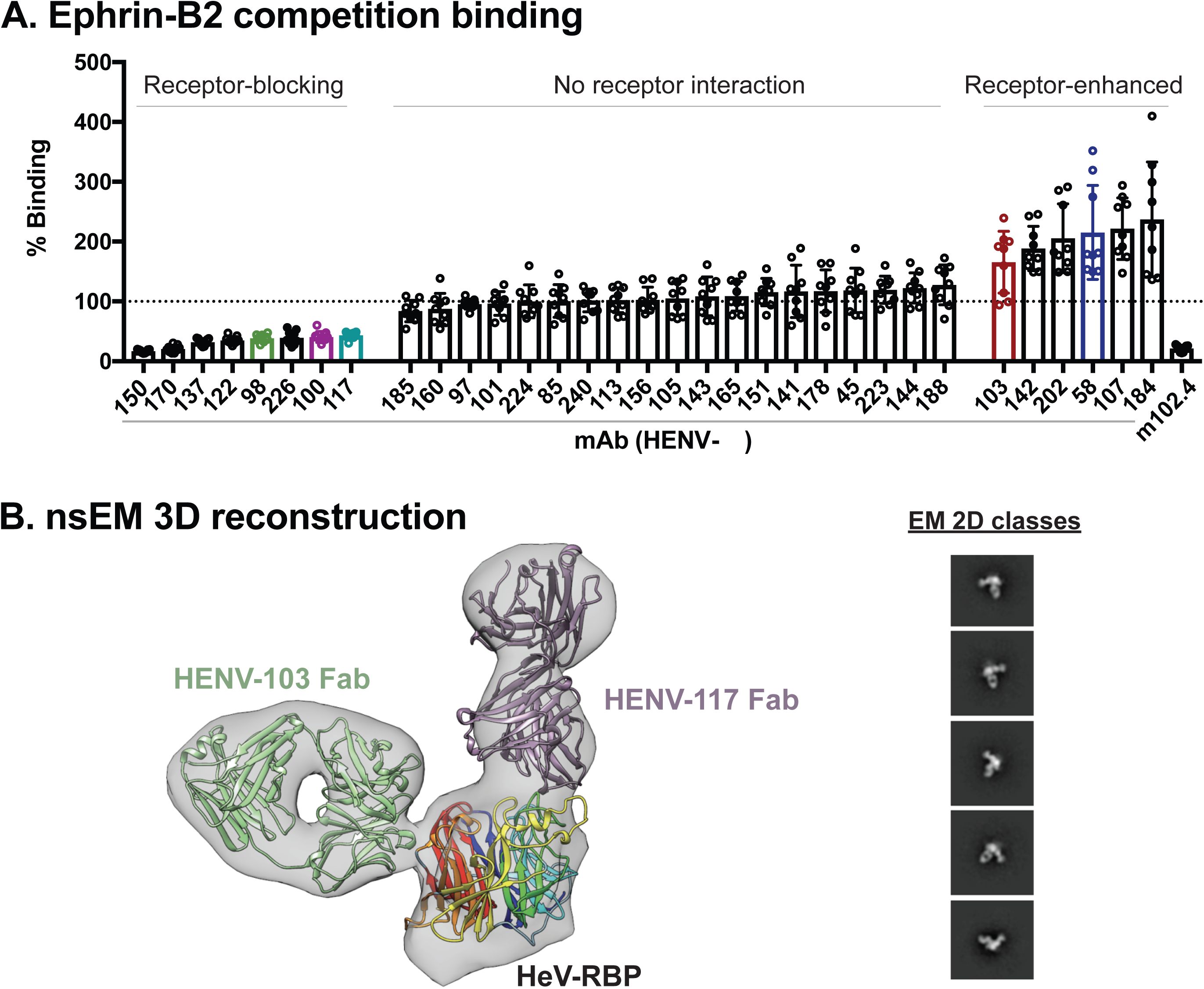
Receptor blocking and structural studies. **A)** Antibody binding to cell-surface-displayed HeV-RBP when ephrin-B2 is bound. Cells transiently transfected with a cDNA encoding the full-length HeV-RBP were incubated with a saturating concentration of recombinantly expressed ephrin-B2. Without washing, cells were incubated with 2 µg/mL antibody, and binding was compared to binding of antibodies in the absence of ephrin-B2. The mAb m102.4 served as a control for receptor competition. Pooled data from 3 independent experiments are shown. **B)** Three-dimensional reconstruction from negative stain electron microscopy of dimeric HeV-RBP full ectodomain bound to HENV-103 Fab and HENV-117-Fab. The EM map is shown in gray, the Fabs are in purple and green, and the RBP head domain is colored by -propeller. 2D classes are shown, with box size of 128 at A/pix of 3.5. β

We also assessed antibodies from all other epitope binning groups for their ability to bind RBP in the presence of ephrin-B2. Antibodies from group F did not bind to surface-displayed HeV-RBP, further suggesting these antibodies cannot access this antigenic site when RBP is in its tetrameric, membrane-anchored form. Group B, C, and E mAbs bound to HeV-RBP with equal signal in the presence or absence of ephrin-B2. Surprisingly, cross-reactive and neutralizing antibodies in group E displayed a “receptor-enhanced” phenotype, in which binding was increased in the presence of ephrin-B2. As HDX experiments suggested this antigenic site lies at the putative interface between protomers within the HeV-RBP dimer, it is likely that receptor binding alters the conformation of HeV-RBP, better exposing this epitope and increasing binding by mAbs to this site. In summary, cross-reactive and neutralizing mAbs displayed either “receptor-blocking” or “receptor-enhanced” phenotypes, suggesting distinct neutralization mechanisms used by antibodies targeting distinct sites.

### Negative stain electron microscopy (nsEM) elucidates structural determinants of recognition by receptor-blocking and receptor-enhanced mAbs

To gain insight into the structural determinants of recognition by “receptor-blocking” and “receptor-enhanced” mAbs, we performed nsEM on HeV-RBP complexed with representative Fabs based on the sequence of HENV-117 (blocking) or HENV-103 (enhanced). Initial studies with HeV-RBP ectodomain (head and stalk domains) purified by size exclusion chromatography showed substantial structural heterogeneity of both dimeric and tetrameric complexes (data not shown). In order to generate more structurally homogeneous antigen suitable for 3D reconstruction, we purified HeV-RBP by gradient fixation ultracentrifugation using a 10 to 30% glycerol gradient containing a linear 0 to 0.1% glutaraldehyde gradient. This method achieved highly pure material, appropriate separation of monomeric, dimeric, and tetrameric species, and structural homogeneity induced by mild glutaraldehyde fixation. Dimeric HeV-RPB was complexed with a molar excess of HENV-117 and HENV-103 and assessed using nsEM.

Both HENV-103 and HENV-117 bind simultaneously to the HeV-RBP, further confirming these mAbs recognize distinct antigenic sites (**Figure 2B**). By docking the crystal model of the head domain bound the ephrin-B2 receptor to the EM map, we observed that HENV-117 mimics the binding position of the receptor, confirming its ability to block receptor attachment (**Figure S3A**). Conversely, HENV-103 approaches the HeV-RBP perpendicular to the receptor binding domain at the putative interface between protomers within the RBP tetramer (**Figure S3B)**. This antigenic site overlaps with previous published mAbs, including HENV-32. Furthermore, modeling suggests that HENV-117 uses a long CDRH3 loop, binding to RBP in a manner similar to the GH loop of ephrin-B2. In summary, HENV-103 and HENV-117 map to distinct antigenic sites by negative stain EM, with HENV-117 mimicking ephrin-B2 binding, while HENV-103 binds at the putative dimeric interface.

### Antibodies provide therapeutic protection in a highly stringent model of Nipah Bangladesh virus challenge in Syrian golden hamsters

Previous studies of murine and human mAbs targeting HeV and/or NiV suggested passive immunization as a potential strategy for therapeutic intervention. To assess therapeutic activity of antibodies in this large panel, we chose 5 candidate mAbs representing groups A (receptor-blocking HENV-98, HENV-100, HENV-117) and D (receptor-enhanced HENV-58, HENV-103) to test in a highly stringent NiV_B_ challenge model in hamsters (Wong et al., 2003). Disease in this model follows a two-stage disease pattern with differing sequelae: an acute respiratory distress syndrome (ARDS)-like respiratory tract component starting at day 3 to 4, and an encephalitic component beginning at days 8 to 12. On day 0, Syrian golden hamsters were challenged intranasally with 5 x 10^6^ PFU NiV_B_. The following day, hamsters were administered a 10 mg/kg dose of antibody by the IP route and monitored for 28 days after challenge. While the hamster administered a vehicle control solution succumbed at day 3, as much as 60% survival was achieved in animals administered either “receptor-blocking” or “receptor-enhanced” mAbs. **(Figure 3A)**. The two most protective mAbs from each class were HENV-117 and HENV-103, for which surviving animals in each treatment group were able to maintain body weight throughout the study **(Figure 3B)**. HENV-117 and HENV-103 were also the two most potent mAbs from groups A and D, suggesting *in vitro* potency by antibodies targeting these sites correlates with *in vivo* efficacy. In summary, receptor-blocking and receptor-enhanced mAbs protect hamsters from NiV_B_ challenge, with HENV-117 and HENV-103 representing the most promising candidates targeting two distinct antigenic sites.

**Figure 3.**
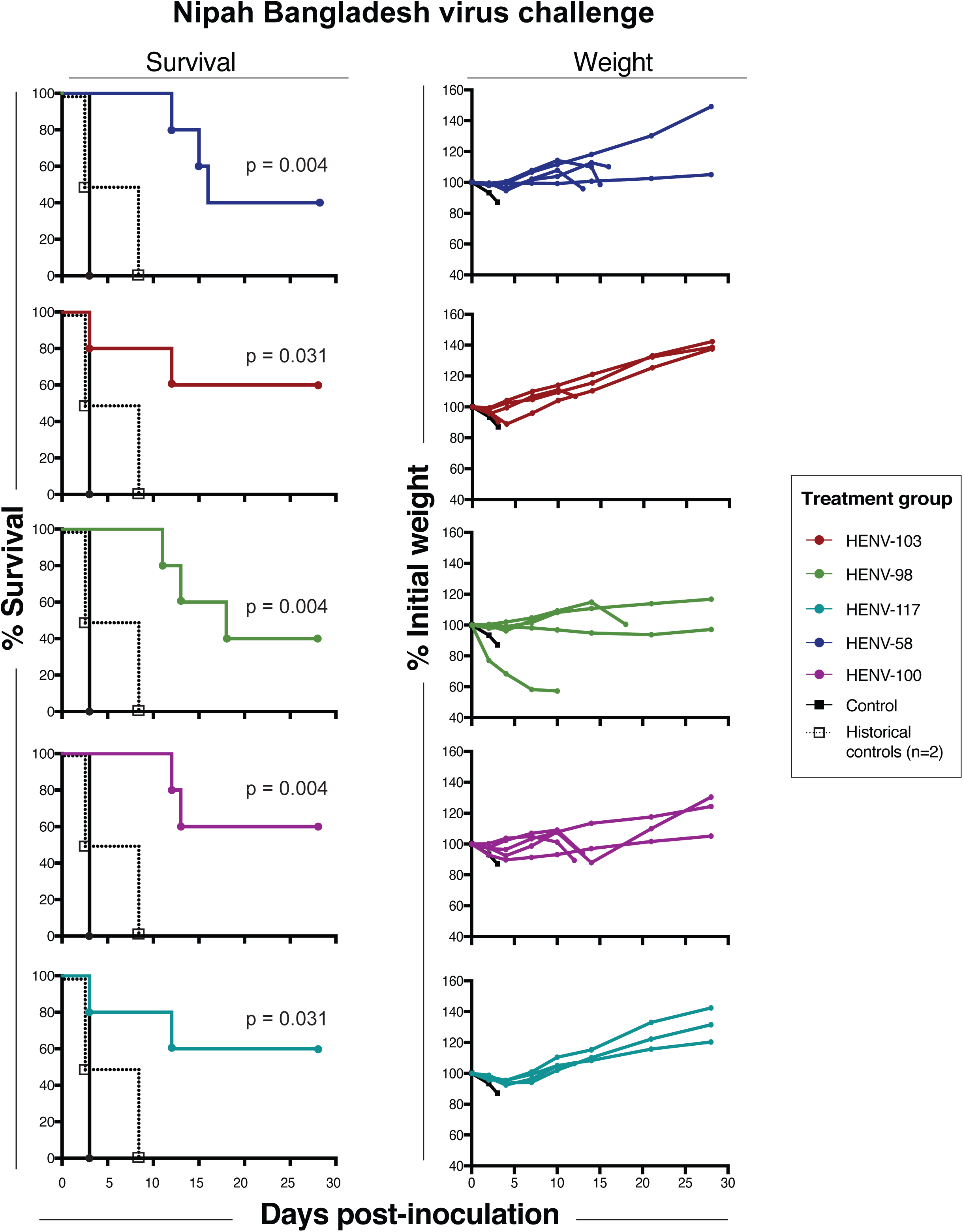
Therapeutic protection by human antibodies in hamster model of Nipah Bangladesh challenge. Survival curves (left) and weight maintenance (right) for hamsters treated with 10 mg/kg antibody (n=5 per group) 24 hours post-inoculation with 5 x 106 PFU NiV Bangladesh by the intranasal route. An untreated control animal (n=1) succumbed to infection 3 days post-inoculation. All weight maintenance charts include control animal in black. Two historical controls are plotted on survival curves and pooled with the experimental control to perform statistical analysis by the long rank Mantel-Cox test.

### HENV-117 and HENV-103 cooperate for binding to HNV-RBP and reveal synergistic virus neutralization activity

RNA viruses, including HeV and NiV, use error-prone RNA-dependent RNA polymerase (RdRP) complexes to achieve genome replication (Welch et al., 2020). While generation of errors can lead to non-viable genomes in some cases, this process also affords viruses the ability to escape from small and large molecule antivirals by introducing amino acid substitutions in the sites recognized by these molecules (Borisevich et al., 2016). This escape pattern is of concern and has been observed in both *in vitro* and *in vivo* studies of diverse RNA viruses, showing that antibody monotherapy approaches against viral pathogens may be susceptible to failure. In order to combat escape, cocktails of antibodies targeting the same or differing antigenic sites offer a higher threshold of protection, with escape becoming statistically highly unlikely. Concurrently, studies of antibody cocktails against Ebola virus, HIV, and more recently SARS-CoV-2, show the potential for synergistic activity by neutralizing antibodies, in which one antibody potentiates the activity of another (Howell et al., 2017; Miglietta et al., 2014; Zost et al., 2020a). With this goal in mind, we sought to determine whether “receptor-blocking” and “receptor-enhanced” mAbs cooperatively bind to and neutralize henipaviruses. We hypothesized that “receptor-blocking” mAbs would mimic the structural rearrangements in HeV-RBP by ephrin-B2, better exposing the “receptor-enhanced” epitope, allowing for synergistic neutralization by combinations of these antibodies. We chose the most potent and protective candidates from each class, HENV-103 and HENV-117, for these studies.

We first tested the ability of HENV-117 to enhance the binding of HENV-103 to cell-surface displayed RBP. Using the surface-display system, we incubated HeV-RBP-transfected cells in saturating concentrations of mAbs that block ephrin-B2 binding. Without washing, we then added serial dilutions of HENV-103 chemically labeled with an Alexa Fluor-647 tag. Cells then were analyzed by flow cytometry to determine if HENV-103 showed increased binding signal across a dilution series in the presence of “receptor-blocking” mAbs. When cells were incubated with HENV-103 only, half maximal binding was achieved at 5,289 ng/mL. When cells were first incubated with saturating concentrations of HENV-117, the EC_50_ of HENV-103 shifted to 350 ng/mL, representing an increase in binding activity of approximately 15-fold (**Figure 4A**).

**Figure 4.**
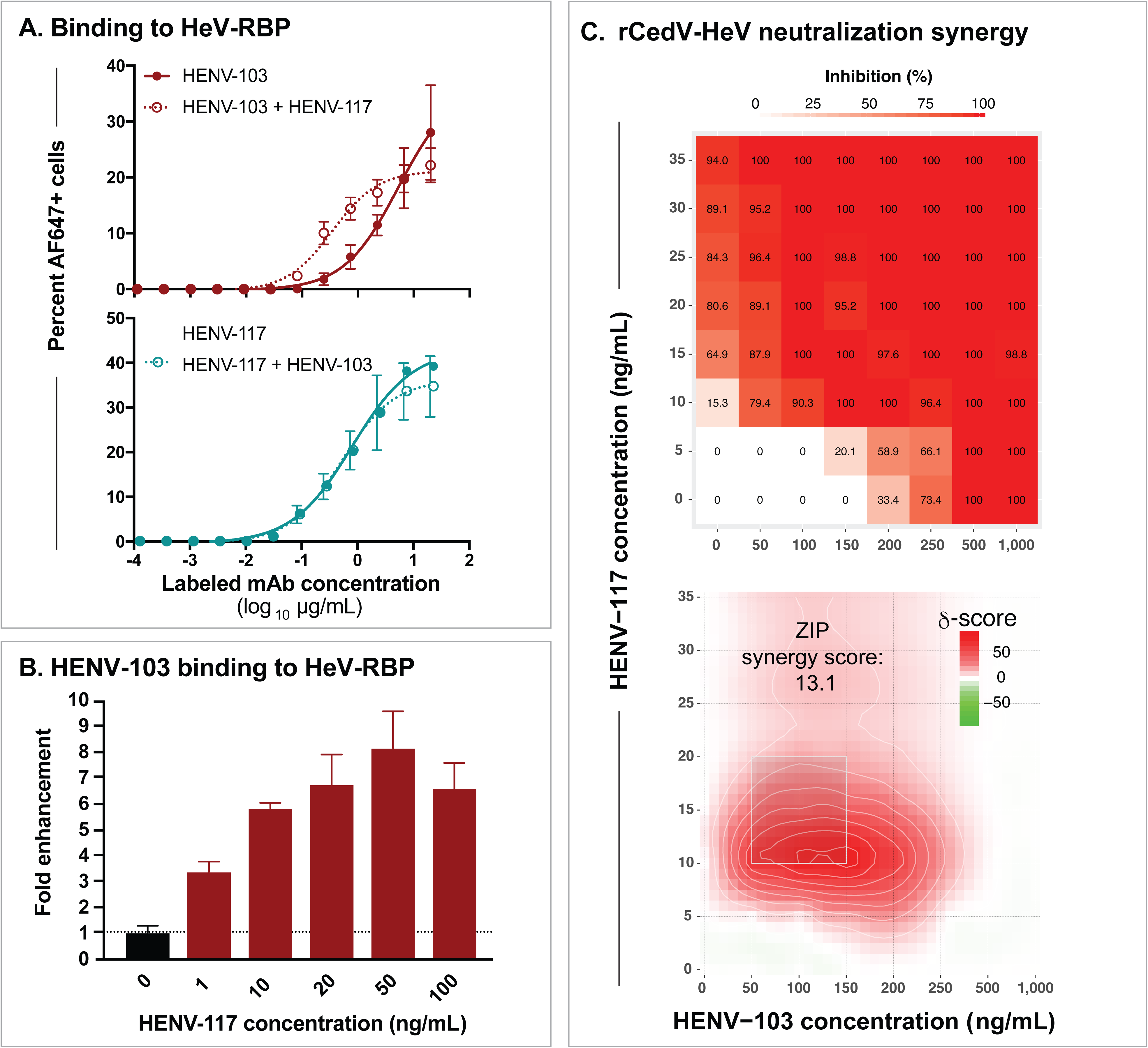
Synergistic binding and neutralization. **A)** Cooperative binding by HENV-103 and HENV-117 to cell-surface-displayed HeV-RBP. Cells expressing HeV-RBP were incubated with unlabeled HENV-103 or HENV-117, followed by addition of a dilution series of Alexa Fluor 647 (AF647) -labeled HENV-103 or HENV-117. Cells were analyzed by flow cytometry and gated for AF647-positive cells. Data were pooled from 3 independent experiments. **B)** Dependence of HENV-117 effective concentration on HENV-103 binding enhancement. Cells were incubated with varying concentrations of unlabeled HENV-117, followed by incubation with AF647-labeled HENV-103 at 0.5 µg/mL, with enhancement calculated by comparing AF647+ cells to HENV-103 binding to HeV-RBP in the absence of HENV-117. Representative data from 3 independent experiments are shown. **C)** Synergistic neutralization of rCedV-HeV by HENV-103 and HENV-117 combinations. Neutralization values at each matrix concentration (top) and calculated synergy scores (bottom) are shown. Serial dilutions of HENV-103 and HENV-117 were mixed with 4,000 PFU rCedV-HeV-GFP for 2 hours, followed by addition to Vero E6 cell monolayers in 96 well plates. Formalin fixed cells were imaged using a CTL S6 analyzer to count GFP+ cells. Neutralization was calculated by comparing treatment to virus- only control wells. Values were imported into SynergyFinder using a Zero Interactions Potency (ZIP) statistical model. Delta scores >10 indicate likely synergy. Two independent experiments were performed, with data from a single representative experiment shown.

Notably, this cooperativity is unidirectional, as HENV-103 did not increase the binding of HENV-117 **(Figure 4A)**. This cooperative phenotype also depends on HENV-117, with increasing HENV-117 concentrations showing increased binding by a constant concentration of HENV-103 **(Figure 4B)**. These data suggest that antibodies that bind the ephrin-B2 binding site on HeV-RBP, such as HENV-117, mimic the conformational changes induced by ephrin-B2 binding, making a semi-cryptic epitope recognized by HENV-103 more accessible.

In order to show that this cooperative binding phenotype is recapitulated functionally, we performed neutralization tests using solutions containing antibody pairs to measure synergistic neutralization potential. In order to perform these multiple comparison neutralization assays quantitatively in BSL-2 facilities, we used a non-pathogenic henipavirus chimerized with the HeV or NiV_B_ glycoproteins. In this system, recombinant Cedar virus (rCedV) was engineered genetically to express the RPB and F from HeV or NiV_B_, as well as a GFP reporter. The resulting rescued, recombinant, chimeric viruses were termed rCedV-NiV_B_ or rCedV-HeV. Here, we used a matrix approach to test antibody pairs for neutralization synergy, in which serial dilutions of HENV-117 and HENV-103 were mixed together in a pairwise matrix, followed by incubation with rCedV-HeV. Virus/mAb mixtures then were added to Vero E6 cell monolayer cultures in 96-well plates. At approximately 22 hours after inoculating cells with virus/antibody mixtures, plates were fixed and GFP+ foci were quantified to enumerate antibody neutralization values. To calculate synergy, neutralization matrix data were uploaded to the open source program “SynergyFinder,” and synergy scores were calculated using the zero interactions potency (ZIP) model (Ianevski et al., 2020). A score >10 suggests synergistic activity. We observed that HENV-103 and HENV-117 gave an overall ZIP score of 13.1, with select physiologically achievable cocktail concentrations achieving synergy scores >20 **(Figure 4C)**. This synergy was also observed when using rCedV-NiV_B_, as well as a VSV psuedotyped with NiV_B_ RBP and F, a platform described in detail previously (**Figure S4A,B**) (Mire et al., 2019). These data together with binding studies show that antibodies from these classes cooperate for binding to RBP and synergistically neutralize chimeric and pseudotyped viruses bearing RBP and F proteins from HeV or NiV_B_, suggesting they will likely function to synergistically neutralize pathogenic henipaviruses.

### Antibody cocktails and derivative bispecific mAbs provide improved therapeutic activity in hamsters

Synergy observed *in vitro* by HENV-103 and HENV-117 against VSV-NiV_B_ and rCedV-NiV_B_ suggested the potential for *in vivo* synergistic protection from NiV_B_ infection. To assess this possibility, we took two separate approaches. In the first approach, we tested HENV-103 and HENV-117 as a cocktail therapy in Syrian golden hamsters. Previously, animals were treated with 10 mg/kg for each individual mAb. In this study, animals were treated with 5 mg/kg HENV-103 and 5 mg/kg HENV-117 at 24 hours after intranasal inoculation with NiV_B_. Using monotherapy, we found that 3 of 5 animals treated with either HENV-103 or HENV-117 survived throughout the study. However, when given in combination, all animals survived and maintained/gained weight for 28 days after infection (**Figure 5B)**. These data show that HENV-103 and HENV-117 provide synergistic protection in hamsters when administered together 24 hours after infection with NiV_B_.

**Figure 5.**
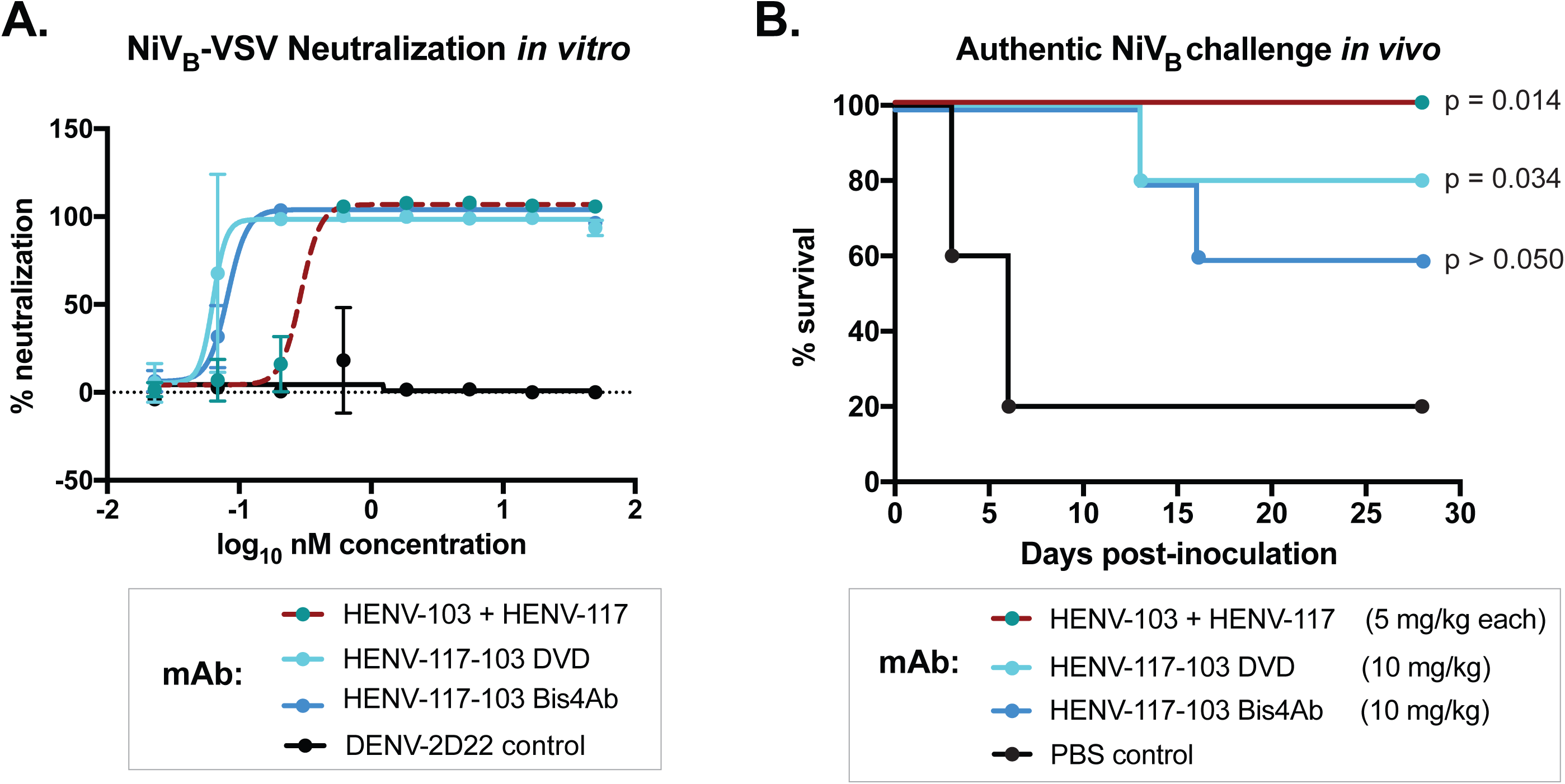
Antibody cocktail and corresponding bispecific antibody therapeutic activity in hamsters. **A)** Neutralization of VSV-NiVB by bispecific antibodies in comparison to equimolar antibody cocktail. Representative data from two independent experiments is shown, each performed in technical triplicate. **B)** Syrian golden hamster challenge studies with HENV-103 and HENV-117 cocktail or corresponding bispecific antibodies. Challenge studies were performed as described above. P values represent statistical significance as determined by Mantel-Cox log rank test. N=5 animals were included in all groups, with control animals treated with PBS at 24 hours post-inoculation.

The second approach used two bispecific antibody platforms. The dual variable domain (DVD) construct bears two heavy and light chain variable domains in each “arm,” with the domains most Fc-distal corresponding to HENV-117 (Wu et al., 2007). A similar construct, termed Bis4Ab, differs from DVD in that the Fc-distal HENV-117 component contains a full Fab fragment, whereas the HENV-103 contains only heavy and light chain variable domains in a Fc-proximal scFv format (Dimasi et al., 2019; Thanabalasuriar et al., 2017). We first tested these constructs *in vitro* against VSV-NiV_B_ and found that both DVD and Bis4Ab constructs strongly neutralized VSV-NiV_B_ with similar potency (**Figure 5A)**. We again tested these in the Syrian golden hamster model of NiV_B_ infection and found that in the DVD group, 4/5 hamster survived, while protection in the Bis4Ab group mirrored that of monotherapy, with 3/5 hamsters surviving. These data suggest that there may be added complexity to using bispecific antibody platforms (whether or not both antigen binding fragments can engage antigen simultaneously, serum half-life in rodents, etc.) and that combined administration of HENV-103 and HENV-117 provides superior *in vivo* protection in comparison to monotherapy. This feature is complemented by the added benefit of further protection from escape mutation.

## Discussion

Recent epidemics of Ebola, 2009 H1N1 influenza, and SARS-CoV-2 viruses highlight the need for the development of countermeasures against emerging viruses with pandemic potential before their occurance. Pathogenic henipaviruses, particularly NiV, are emerging and highly pathogenic agents with confirmed human-to-human transmission for which licensed treatments or vaccines for human use do not yet (Amaya and Broder, 2020). In this study, we isolated a panel of mAbs specific for the henipavirus RBP glycoprotein from an individual with prior occupation-related exposure to equine HeV-RBP subunit vaccine. Competition-binding and HDX-MS studies identified at least six distinct antigenic sites recognized by these mAbs. Flow cytometric studies with surface-displayed HeV-RBP showed that potently neutralizing, cross-reactive antibodies either a) blocked HeV-RBP binding to ephrin-B2, or b) showed enhanced binding in the presence of ephrin-B2. Antibodies that block receptor binding also induced the “receptor-enhanced” phenotype, showing that antibodies to these two classes cooperate for binding to HNV-RBP. In addition, these mAbs also showed synergy in neutralization of rCedV-HeV particles. As monotherapy, “receptor-blocking” and “receptor-enhanced” antibodies provided modest protection in a stringent, highly lethal NiV_B_ challenge model in Syrian golden hamsters. In combination, these antibodies provided complete therapeutic protection in the same model of infection.

A significant concern when using antibodies as therapeutics against emerging infectious diseases due to RNA viruses is the potential for viral ‘mutational escape’ within an infected host or immune evasion by divergent viral variants. Escape from antibody-mediated neutralization has been documented even with ultrapotent mAbs targeting conserved epitopes on viral glycoproteins (Greaney et al., 2020). Using a cocktail of mAbs provides resistance against escape, with the potential added benefit of synergistic antiviral potency, allowing for lower dosing. The potential for spillover of divergent variants of bat-borne paramyxoviruses (henipaviruses and rubulaviruses) is consistent with the inherent propensity of RNA viruses for rapid evolution. Furthermore, flying foxes serve as ideal reservoir hosts because of their dense community roosting patterns and relative resistance to paramyxoviral disease (Baker et al., 2012; Barr et al., 2015; Drexler et al., 2012; Luis et al., 2015; Peel et al., 2019; Sasaki et al., 2012; Vidgen et al., 2015). The discovery of protective antibodies highlighted here, specifically HENV-103 and HENV-117, offer the opportunity to construct a cocktail of antibodies with most-desired protective properties including against mutation escape and spillover variant viruses. Concurrently, a bispecific antibody with activity of both HENV-103 and HENV-117 is an attractive therapeutic option that endows a single therapeutic molecule with the synergistic potency of two individual mAbs. While we showed that two antibodies targeting RBP offer a synergistic benefit, the possibility exists that having antibodies targeting both RBP and F may provide also be of benefit. Recently, highly potent and protective anti-F antibodies have been described and may offer an ideal partner to HENV-103, HENV-117, or both as a triple antibody cocktail (Dang et al., 2019; Mire et al., 2020).

As with other paramyxoviruses, humans likely elicit highly functional antibodies against the henipavirus F glycoprotein (Merz et al., 1980). This concept is highlighted by palivizumab, an anti-F antibody used as a prophylaxis for premature infants to protect from infection by respiratory syncytial virus (Meissner et al., 1999). Although, as highlighted above, protective anti-F mAbs have been isolated, these have been uniformly of murine origin. The full antigenic landscape of the henipavirus F protein may suggest new sites of vulnerability to neutralization by mAbs and could guide the rational design of henipavirus vaccines. This opportunity is especially important considering the geographical range of henipaviruses, and the fact that a previously undescribed virus from this genus may emerge to cause a pandemic. Having knowledge of the determinants of neutralization for both RBP and F will allow for quick mobilization of platform technologies to develop vaccines, similar to what we have seen in the response to the SARS-CoV-2 pandemic (Zost et al., 2020b).

Wild-type mice are understood to be refractory to infection with henipaviruses. On the contrary, the Syrian golden hamster model of henipavirus infection has been demonstrated to recapitulate the most salient features of human disease, including the biphasic pulmonary disease followed by catastrophic neurological events (Rockx et al., 2011), making it an ideal model to screen vaccines and therapeutics. However, hamsters require an orders of magnitude higher challenge dose in order to achieve lethal disease compared to ferrets or African Green monkeys (AGM). This requirement likely contributes to a much faster disease progression and a concievably shorter therapeutic window. Ferrets, and optimally, AGM are likely superior animal models to further preclinical development of these promising antibody candidates. This possibility is especially true of the AGM model of NiV_B_ infection, in which the therapeutic window for use of antibody therapies (treatment at days 3 to 5) is shorter than that of HeV and NiV_M_ (treatment at days 5 to 7) (Mire et al., 2016). Studies in these models might further elucidate if HENV-103 and HENV-117 are improved compared to previously described antibodies (Playford et al., 2020).

Here, and in previous studies, functional anti-henipavirus RBP-specific mAbs from multiple species have been isolated. These antibodies uniformly recognize the head domain of RBP, suggesting this region is likely the most immunogenic domain of the RBP. Multiple studies interrogating the function of RBP, and its role in triggering the F protein to undergo significant conformational rearrangements, have pointed to the RBP stalk domain as playing a significant role in viral fusogenicity. Specifically, Aguilar *et al*. have shown that the C-terminal portion of the NiV stalk domain can trigger fusion of membranes in the absence of a head domain (Liu et al., 2013). While it is likely that the stalk domain, which is partially obstructed by the head domain, is immunogenically subdominant, it is possible that rare, circulating memory B cells harboring antibodies targeting this domain exist. Future studies interrogating the antibody response to these viruses also may shed light on the role of mAbs targeting the stalk domain of HNV-RBP, and whether these antibodies have the potential to prevent viral and host membrane fusion.

## SUPPLEMENTAL INFORMATION

Supplemental Information including supplemental experimental procedures, figures, and tables can be found with this article online.

## ACKNOWLEDGEMENTS

We thank Chris Gainza and Rachel Nargi from Vanderbilt in the Crowe laboratory for technical support with antibody production and purification, and Joseph Reidy, Andrew Trivette, and Robert Carnahan for intellectual contributions to antibody production and sequencing, and Ian Setliff and Ivelin Georgiev for statistical support. We also thank Mathew Hyde and the UTMB Animal Resource Center for their assistance with animal procedures. This work was supported by a grant from the National Institutes of Health (NIH) U19 AI142764 and departmental funds from UTMB to Thomas Geisbert and UC7AI094660 for BSL-4 operations support of the Galveston National Laboratory. The work of MPD was supported by NIH fellowship grant F31 AI152332. The project described was supported by CTSA award No. UL1 TR002243 from the National Center for Advancing Translational Sciences (NCATS). The contents of this publication are solely the responsibility of the authors and do not necessarily represent the official views of NIAID or NIH.

## AUTHOR CONTRIBUTIONS

Conceptualization M.P.D. and J.E.C.; Methodology M.P.D., N.K., V.B., I.S., I.G., K.L.S., E.A.K., B.R.W.; Investigation M.P.D., N.K., V.B., E.B., M.A., M.N., E.A., R.B., J.D., I.S., K.L.S., L.Z., E.A.K., Z.A.B., B.R.W., T.W.G., R.W.C., J.E.C.; Resources E.J.A., C.C.B., T.W.G., J.E.C.; Writing – Original Draft M.P.D., J.E.C., Writing – Review & Editing all authors; Supervision C.C.B., T.W.G., J.E.C.; Project Administration T.W.G., J.E.C.; Funding Acquisition M.P.D., L.Z., T.W.G., J.E.C., C.C.B.

## Declaration of interests

J.E.C. has served as a consultant for Lilly and Luna Biologics, is a member of the Scientific Advisory Boards of CompuVax and Meissa Vaccines and is Founder of IDBiologics. The Crowe laboratory at Vanderbilt University Medical Center has received unrelated sponsored research agreements from IDBiologics and AstraZeneca. E.A.K., Z.A.B., and B.R.W. are employees and shareholders of Mapp. L.Z. is an employee, shareholder and co-owner of Mapp. All other authors declare no competing interests.

**Figure S1 (related to figure 1). ELISA binding curves for representative antibodies from group A (receptor-blocking), group D (receptor-enhanced), or control antibodies.** Values were fit to a non-linear regression model in GraphPad to generate curves following log transformation of antibody concentrations. Data representative of three independent experiments performed in technical duplicate are shown.

**Table S1 (related to Figure 1). Sequence features of 5 potently neutralizing, cross reactive antibodies chosen for *in vivo* studies.** Heavy and light chain sequences were analyzed using IgBLAST to identify V(D)J pairings for each mAb shown.

**Figure S2 (related to Figure 1). Pearson correlation analysis of surface plasmon resonance competition binding.**

**Figure S3 (related to Figure 2). Prediction of antigenic sites recognized by HENV-103 and HENV-117 by negative stain EM.** A) The RBP head domain with a model of HENV-117 scFv (gray) overlapping with ephrin-B2 receptor electron density (dot surface PRB: 9PDL). B) The RBP head domain with a model of HENV-103 Fv (blue) from top (right, looking down on ephrin-B2 binding face) and side (left) view. Black line denoted the putative dimerisation interface

**Figure S4 (related to Figure 3). Synergistic neutralization of rCedV-NiV_B_ and VSV-NiV_B_ by HENV-103 and HENV-117**. Synergy plots using the zero interactions potency (ZIP) model generated by SynergyFinder for neutralization of A) rCedV-NiV_B_ or B) VSV-NiV_B_. Representative data from two (A) or three (B) independent experiments shown.

## Methods

### Generation of hmAbs

Peripheral blood mononuclear cells (PBMCs) from a human subject were isolated from whole blood and transformed using Epstein-Barr virus (EBV), as previously described (Crowe, 2017). Briefly, transformed B cells were expanded and co-cultured with irradiated human PBMCs in 96-well plates. Cell supernatants were screened by ELISA using recombinant HeV-RBP or NiV-RBP head domain proteins. Wells with positive reactivity then were fused to a human-mouse heteromyeloma cell line (HMMA 2.5) and plated by limiting dilution in 384-well plates. The resulting hybridomas were cloned as single cells by fluorescence-activated cell sorting (FACS) to produce clonal hybridoma cell lines. These clonal hybridoma cells were cultured in T-225 flasks containing serum-free medium, and mAb was purified from spent medium by affinity chromatography on an ÄKTA^TM^ pure Fast Protein Liquid Chromatography (FPLC) instrument (GE Healthcare).

### Generation of bispecific mAbs

Bispecific mAbs that combined the antigen binding domains of HENV-117 and HENV-103 into a single molecule were designed, expressed, and purified as follows. The heavy chain of the HENV-117-103 DVD combines the heavy chain variable domains of first HENV-117, then HENV-103, each separated by a flexible linker, and then followed by the IgG1 human constant heavy chain domains. Similarly, the light chain of the HENV-117-103 DVD includes the light chain variable domains of both HENV-117 and HENV-103, separated by a flexible linker and then followed by a single human kappa light chain constant domain which naturally pairs with the corresponding DVD heavy chain. The HENV-117-103 Bis4Ab was constructed by inserting a HENV-103 single-chain variable fragment (scFv) between CH_1_ and CH_2_ of the HENV-117 heavy chain. The HENV-103 scFv in the bis4Ab format contains a poly glycine-serine linker between its variable domains, and the scFv unit is also flanked by poly glycine-serine linkers.

The modified heavy chain is then paired with the standard HENV-117 light chain for expression and purification (Dimasi et al., 2019). The heavy and light chains of the HENV-117-103 DVD and the HENV-117-103 Bis4Ab were cloned into pcDNA3 expression vectors. For each of the bispecific mAbs, the corresponding heavy and light chain plasmids were chemically co-transfected into ExpiCHO cells (Gibco) and transiently expressed for 9 days. The supernatant was then clarified by centrifugation and filtration, prior to loading onto a MabSelect SuRe Protein A (GE Healthcare) affinity chromatography column using an ÄKTA^TM^ Fast Protein Liquid Chromatography (FPLC) instrument (GE Healthcare). The column was washed with 1X PBS, and the mAbs were eluted with IgG Elution Buffer (Pierce). Following neutralization with 1 M Tris pH 8.0 to pH ∼7, the eluates were concentrated to 5 mg/ml in an Amicon 30K MWCO centrifugal filter (Millipore), and then sterile-filtered using a 0.22 µM syringe filter (Millex-GP).

### HeV and NiV viruses

NiV number 1999011924 was obtained from a patient from the 1999 outbreak in Malaysia. The isolate of NiV_B_ used was 200401066 and was obtained from a fatal human case during the outbreak in Rajbari, Bangladesh in 2004 and passaged on Vero E6 cell monolayer cultures three times. HeV was obtained from a patient from the 1994 outbreak in Australia. All viruses were kindly provided by Dr. Thomas Ksiazek, UTMB. Each virus was propagated on Vero E6 cells in Eagle’s minimal essential medium supplemented with 10% fetal calf serum. The NiV_M_, NiV_B_ and HeV challenge virus stocks were assessed for the presence of endotoxin using The Endosafe-Portable Test System (PTS) (Charles River Laboratories, Wilmington, MA). Each virus preparation was diluted 1:10 in Limulus Amebocyte Lysate (LAL) Reagent Water per the manufacturer’s instructions, and endotoxin levels were tested in LAL Endosafe-PTS cartridges as directed by the manufacturer. Each preparation was found to be below detectable limits, whereas positive controls showed that the tests were valid. All experiments involving infectious henipaviruses were carried out at the UTMB Galveston National Laboratory under biosafety level 4 conditions.

### Neutralization assays

The virus neutralizing activity concentrations were determined for NiV_M_, NiV_B,_ and HeV using a plaque reduction assay. Briefly, antibodies were diluted two-fold from g/mL to extinction and incubated with a target of ∼100 plaque-forming units (pfu) of NiV_M_, NiV_B,_ and HeV45 min at 37 °C. Virus and antibody mixtures then were added to individual wells of six-well plates of Vero 76 cell monolayer cultures. Plates were fixed and stained with neutral red two days after infection, and plaques were counted 24 h after staining. Neutralization potency was calculated based on pfu for each virus in the well without antibody. The neutralization experiments were performed in triplicate, with independent virus preparations and duplicate readings for each replicate. Mean half-maximal inhibitory concentration (IC_50_) values were calculated as previously described (Ferrara and Temperton, 2018).

### Surface plasmon resonance (SPR) epitope binning

A continuous flow micro-spotter (CFM) instrument (Carterra) was used to generate antibody-coated SPR sensor chips (Xantec) (Abdiche et al., 2014). Briefly, mAbs were diluted to 10 µg/mL in sodium acetate pH 4.5 in a 96-well round bottom plate. A mirroring 96-well plate containing activation buffer (EDC and sulfo-NHS in 10 mM MES pH 5.5) was used first to activate the gold-plated surface of the sensor chip, followed by association of antibodies. The coated chip then was moved to an IBIS-MX96 microarray-based surface plasmon resonance imager (Carterra), where it was quenched with 1 M ethanolamine to prevent further coupling of proteins. To bin antibodies, 100 mM HeV-RBP head domain was flowed over the coated sensor chip. One-by-one, antibodies diluted to 10 µg/mL were tested for their ability to associate with antigen captured on the sensor chip. Carterra Epitope software was used to analyze data and construct competition-binding grids.

### Hydrogen-deuterium exchange mass spectrometry (HDX-MS) of Fab-HeV-RBP complexes

HDX-MS was performed as previously reported (Bennett et al., 2019). Briefly, antigen (HeV-RBP) and selected mAbs were prepared individually or in complex at a protein-concentration of 10 pmol/µL in 1× PBS pH 7.4 and incubated for 2 h at 0 °C. Deuterium labeling was performed by a 20-fold dilution of 3 µL sample in PBS pH 7.4 in D_2_O and incubation at 20 °C for 0 s, 100 s, and 1,000 s. The reaction was quenched by a 2-fold dilution in 1× PBS, 4 M guanidinium/HCl, 100 mM tris(2-carboxyethyl)phosphine to a final pH of 2.3 at 0 °C. Samples were injected immediately into a nano-ACQUITY UPLC system controlled by an HDX manager (Waters Corporation, Milford, MA, USA). Online pepsin digestion was performed at 15 °C, 10,000 psi at a flow of 100 µL/min of 0.1% formic acid in H_2_O using an immobilized-pepsin column. A Waters VanGuard™ BEH C18 1.7 μm guard column was used to trap peptides at 0 °C for 6 min before separation on a Waters ACQUITY UPLC BEH C18 1.7 µm, 1 mm × 100 mm column at a flow of 40 µL/min at 0 °C with a 6 min gradient of 5 to 35% acetonitrile, 0.1% formic acid in H_2_O. UPLC effluent was directed into a Waters Xevo G2-XS with electrospray ionization and lock-mass acquisition (human Glu-1-Fibrinopeptide B peptide, m/z=785.8427) for peptide analysis in MS^E^-mode. The capillary was set to 2.8 kV, source-temperature to 80 °C, desolvation temperature to 175 °C, desolvation gas to 400 L/h and the instrument was scanned over a m/z-range of 50 to 2000. All experiments were carried out in triplicate. Data analysis was accomplished using Waters ProteinLynx Global Server 3.0.3 software (non-specific protease, min fragment ion matches per peptide of three, FDR 4% and oxidation of methionine as a variable modification) for peptide identification and DynamX 3.0 software (minimum intensity of 500, minimum products 3, minimum products per amino acid 0.3 and a mass error < 15 ppm) for deuterium uptake calculations. Results are reported as an average of triplicate analyses.

### Generation of VSV pseudotyped viruses bearing NiV_B_ glycoproteins

Recombinant VSVs containing genomic inserts for expression of NiV_B_ G and F proteins were kindly provided by Chad Mire and generated as previously described (Mire et al., 2019). Stocks of each rVSV were propagated and titrated on VSV-G transfected BHK-21 (WI-2), with viral titers determined by counting GFP+ cells using a CTL S6 Analyzer instrument (company/). To generate virus bearing both G and F glycoproteins, cells were inoculated with each VSV at MOI=5 and incubated for 48 hours. Cell supernatants were clarified by centrifugation. Resulting VSV-NiV_B_ was titrated on Vero cell monolayers using an xCELLigence instrument to determine the lowest virus concentration that would induce CPE as measured by cell impedance.

### Cooperative binding of antibodies to antigen displayed on the surface of cells

A construct containing cDNA encoding full-length HeV-RBP protein was transfected using polyethylenimine into 293F cells, and cells were cultured at 37 °C in 5% CO_2_ for 3 days. Cells subsequently were plated at 50,000 cells/well in V-bottom 96-well plates, washed, and incubated with either 20 µg/mL primary mAb in 30 µL or FACS buffer alone for 30 minutes at 4 °C. Without washing, 30 µL serially diluted mAb labeled with Alexa Fluor 647 dye (ThermoFisher) was added to wells and incubated for 30 minutes at 4 °C. Cells were washed and resuspended in FACS buffer and analyzed using an iQue Plus flow cytometer (Intellicyt). Dead cells were excluded from analysis by fluorescent staining with 4 ,6-diamidino-2-phenylindole (DAPI).

### Negative stain electron microscopy

For electron microscopy imaging of HeV-RBP protein and Fabs complex, we expressed the HeV-RBP full ectodomain (head domain with intact stalk domain) with a C-terminal polyhistidine tag. Expressed protein was isolated by metal affinity chromatography on HisTrap Excel columns (GE Healthcare), followed by GraFix methods using a 10% to 30% glycerol gradient and 0 to 0.1% glutaraldehyde gradient (Stark, 2010). Glutaraldehyde was quenched with 1 M Tris-Cl after fractionation. 200 µL fractions were analyzed by SDS-PAGE, with fractions corresponding to monomeric, dimeric, and tetrameric species pooled. Protein was then buffer exchanged into 50 mM Tris-Cl pH 7.5 containing 140 mM NaCl. Fabs corresponding to HENV-103 and HENV-117 were expressed and purified as previously described. Protein complexes were generated by incubation of HeV-RBP_ecto_ dimer and the two Fab in a 1:5:5 molar ratio L of the sample at ∼10 to 15 μg/mL was applied to a glow-discharged grid with continuous carbon film on 400 square mesh copper electron microscopy grids (Electron Microscopy Sciences). Grids were stained with 0.75% uranyl formate (Ohi et al., 2004). Images were recorded on a Gatan US4000 4k × 4k CCD camera using an FEI TF20 (TFS) transmission electron microscope operated at 200 keV and control with SerialEM. All images were taken at 62,000× magnification with a pixel size of 1.757 Å per pixel in low-dose mode at a defocus of 1.5 to 1.8 μm. The total dose for the micrographs was ∼35 e− per Å^2^. Image processing was performed using the cryoSPARC software package. Images were imported, CTF-estimated and particles were picked automatically. The particles were extracted with a box size of 256 pixels and binned to 128 pixels (pixel size of 3.514 A/pix) and 2D class averages were performed to achieve clean datasets. Classes were further classified (2D) to separated different complex variant and classes having the 2 Fab on one RBP domain were selected. *Ab-initio* was used to generate initial 3D volume that was further refined with a mask over one RBP-Fabs complex. The final refine volume has a resolution of ∼15Å. Model docking to the EM map was done in Chimera (Pettersen et al., 2004). For the RBP head domain PDB: 6PDL was used and for the Fab PDB:12E8 or the prediction model of the Fv that was generated by SAbPred tool was used (Dunbar et al., 2016). The 3D EM map has been deposited into EMDB (EMDB XXX). Chimera software was used to make all structural figures.

### Neutralization synergy of VSV-NiV_B_

VSV-NiV_B_ pseudotype viruses were generated as described above. In 96-well plates, serial dilutions of “receptor-blocking” and “receptor-enhanced” mAbs were mixed in a matrix arrangement, followed by addition of equal volume of VSV-NiV_B_ diluted 1:500 in DMEM. Mixtures were incubated for 1 hour at 37 °C prior to addition to Vero cell monolayers in xCELLigence 96-well E-plates containing 10,000 cells/well. Cells were incubated with virus and antibody for 1 hour at 37 °C, followed by addition of DMEM + 5% FBS to wells. Plates were placed back on the xCELLigence instrument for reading of cell impedance every 15 minutes for 72 hours. Neutralization was determined by comparing cell index of treated wells vs. untreated wells at a single time point (values output by xCELLigence instrument software). Neutralization values then were imported into SynergyFinder software (Ianevski et al., 2020), with delta scores calculated using the zero interactions potency (ZIP) synergy model.

### Neutralization synergy of rCedV chimeric viruses

Recombinant Cedar virus (rCedV) chimeras displaying RBP and F proteins of HeV or NiV_B_ were generated and validated as described elsewhere. Black-walled 96-well plates (Corning Life Sciences; NY, USA) were coated with 20,000 cells/well Vero E6 cells in DMEM + 10% Cosmic calf serum and incubated overnight. Approximately 24 hours later, HENV-103 and HENV-117 were diluted to indicated concentrations and incubated 1:1 with 4,000 PFU/well rCedV-HeV-GFP or rCedV-NiV_B_-GFP and incubated for 2 hours at 37 °C. Following incubation, 90 µL virus/antibody mixtures were added to aspirated cell monolayers and were incubated at 37 °C for 22 hours. Medium containing virus/antibody mixtures was aspirated, and cells were fixed with 100 µL/well 4% formalin for 20 minutes at room temperature. After aspiration, cell monolayers were gently washed 4x with DI water. Formalin-fixed plates were then scanned using the CTL S6 analyzer (Cellular Technology Limited; OH, USA). Fluorescent foci were counted using the CTL Basic Count™ feature and S6 software. Percent neutralization was calculated by normalizing counts to a virus only control. Matrices were then imported into SynergyFinder and analyzed as described before.

### Antibody therapy in Syrian golden hamster model of Nipah Bangladesh

3 to 5 week-old Syrian golden hamsters were inoculated with 5 x 10^6^ PFU Nipah Bangladesh (passage 3) via the intranasal route. At 24 hours post challenge, 5 animals per group were treated with 10 mg/kg antibody by intraperitoneal administration. Animals were monitored for 28 days for changes in weight, temperature, and clinical appearance. Animals were humanely euthanized at the experimental endpoint. A single untreated animal served as a control in each study highlighted.

