## Supplementary figures and images for "Functional cooperativity mediated by rationally selected combinations of human monoclonal antibodies targeting the henipavirus receptor binding protein"

### Supplemental Figures

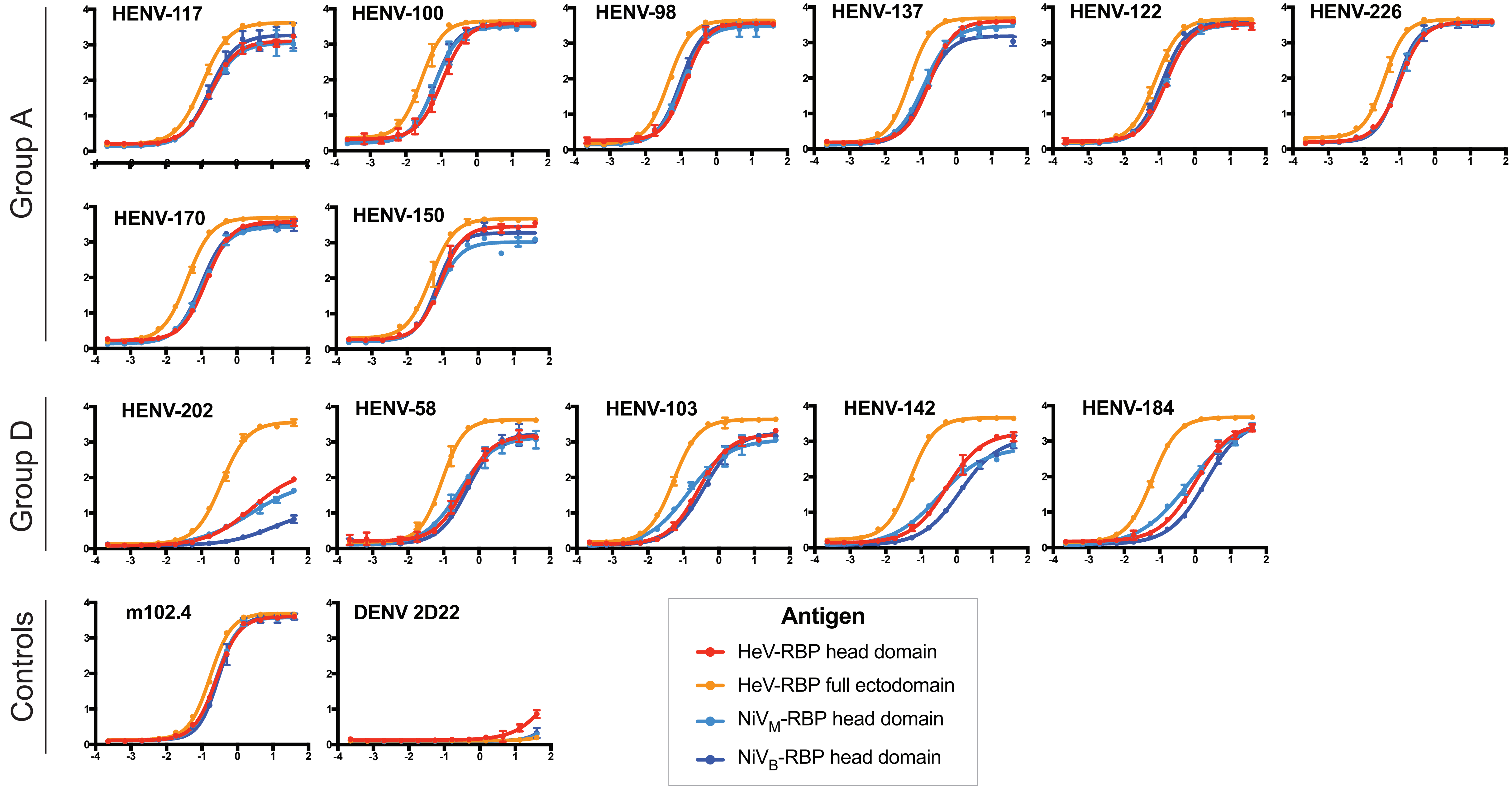

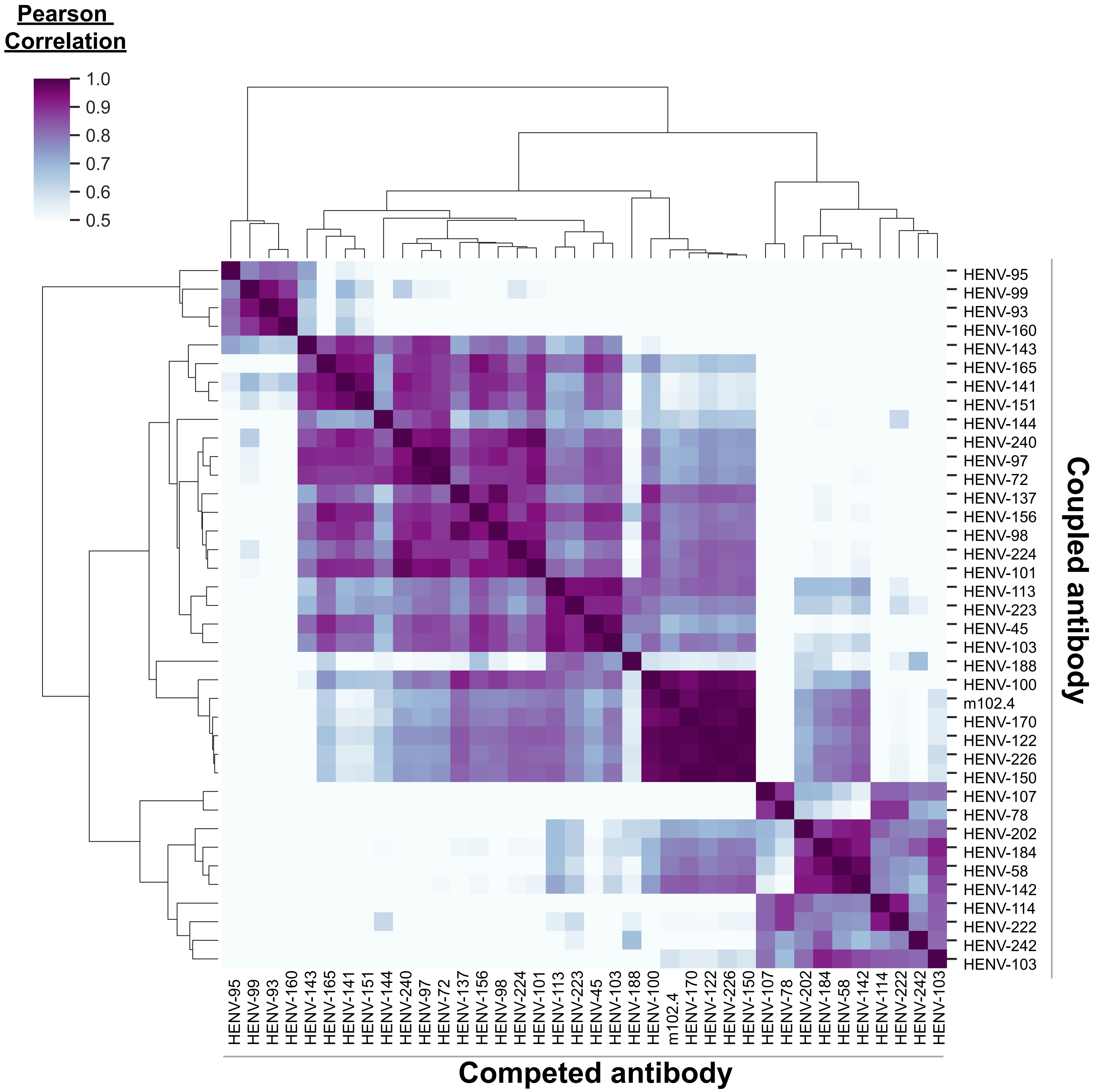

A. Overlay of HENV-117 Fab and ephrin-B2

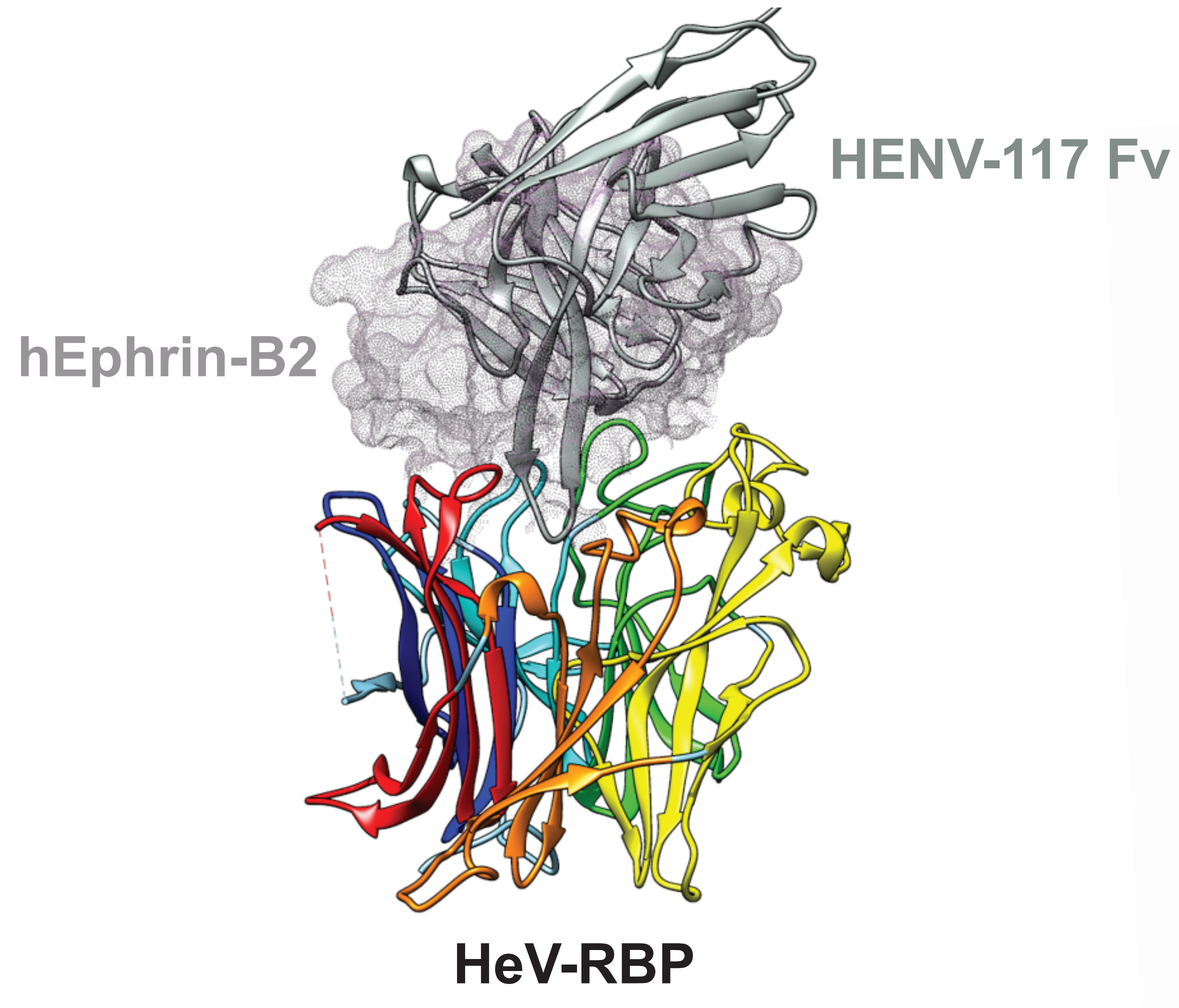


B. Prediction of HENV-103 antigenic site

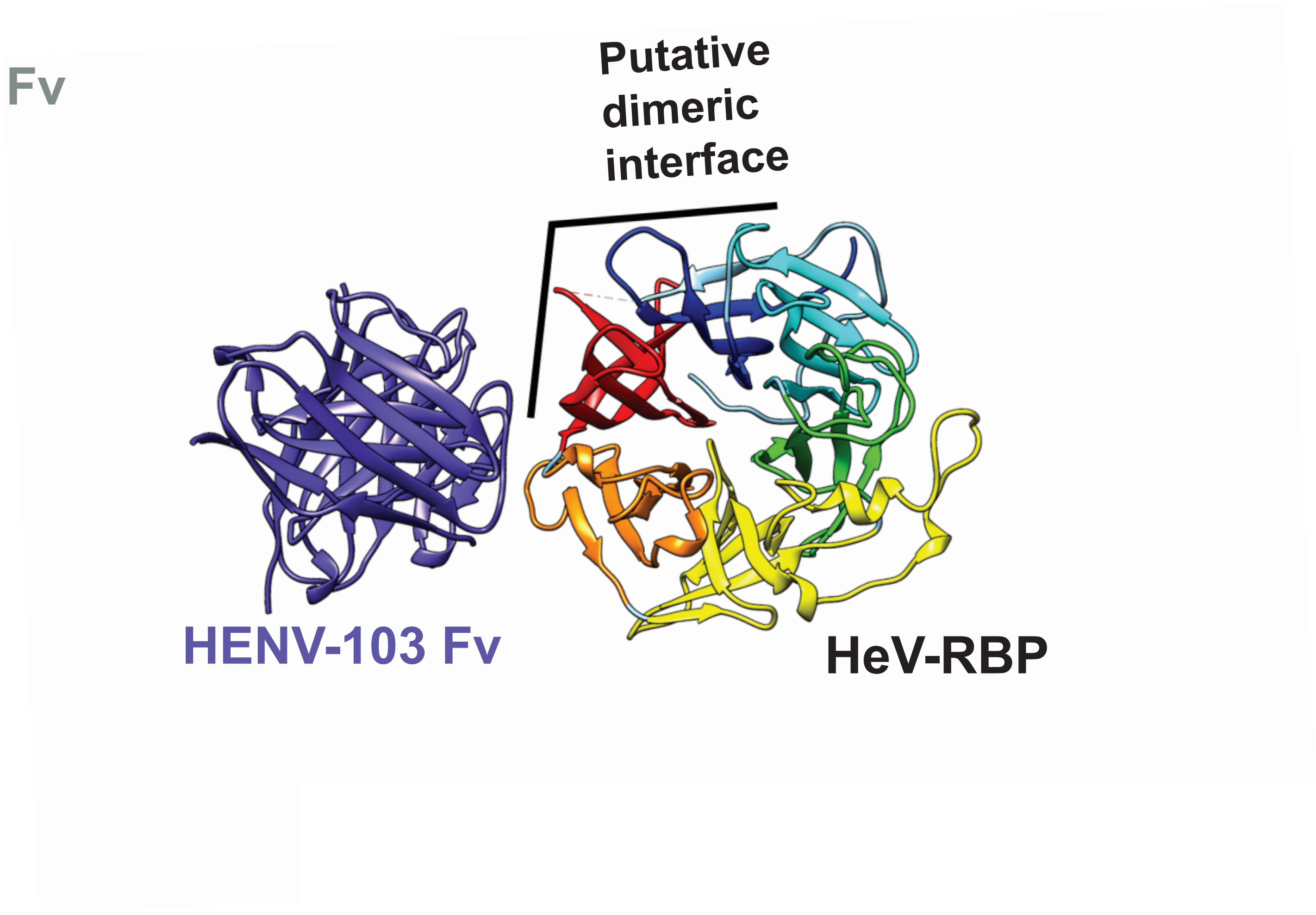

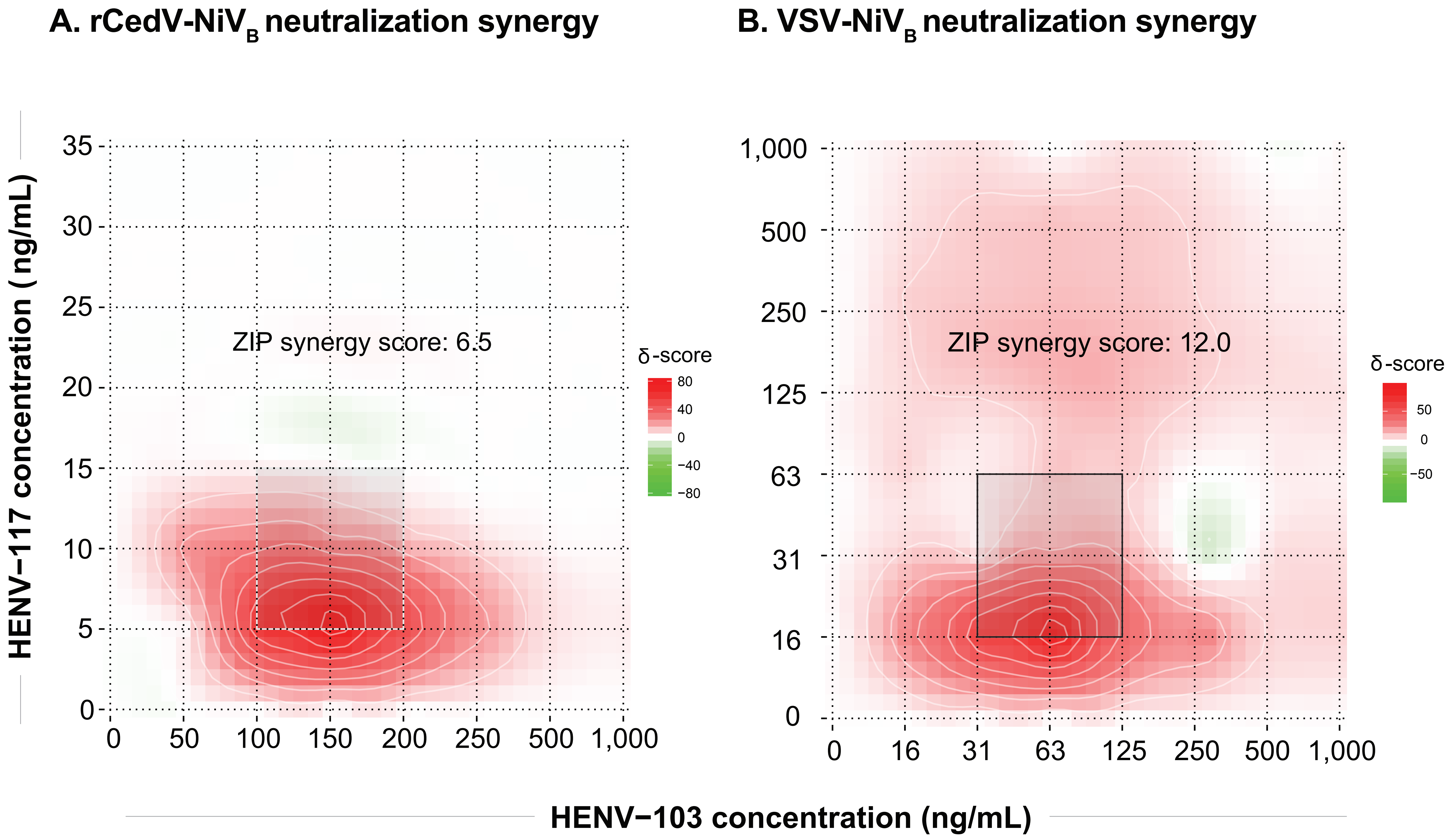
